# PoSTcode: Probabilistic image-based spatial transcriptomics decoder

**DOI:** 10.1101/2021.10.12.464086

**Authors:** Milana Gataric, Jun Sung Park, Tong Li, Vasyl Vaskivskyi, Jessica Svedlund, Carina Strell, Kenny Roberts, Mats Nilsson, Lucy R. Yates, Omer Bayraktar, Moritz Gerstung

## Abstract

Realising the full potential of novel image-based spatial transcriptomic (IST) technologies requires robust and accurate algorithms for decoding the hundreds of thousand fluorescent signals each derived from single molecules of mRNA. In this paper, we introduce PoSTcode, a probabilistic method for transcript decoding from cyclic multi-channel images, whose effectiveness is demonstrated on multiple large-scale datasets generated using different versions of the *in situ* sequencing protocols. PoSTcode is based on a re-parametrised matrix-variate Gaussian mixture model designed to account for correlated noise across fluorescence channels and imaging cycles. PoSTcode is shown to recover up to 50% more confidently decoded molecules while simultaneously decreasing transcript mislabeling when compared to existing decoding techniques. In addition, we demonstrate its increased stability to various types of noise and tuning parameters, which makes this new approach reliable and easy to use in practice. Lastly, we show that PoSTcode produces fewer doublet signals compared to a pixel-based decoding algorithm.

## Introduction

A series of powerful spatial transcriptomics (ST) protocols have been developed in the last decade, which can measure and localise gene expression in its tissue context, thereby providing a novel way to augment genomics with spatial information and thus better understand how tissue function is established and cellular interactions mediated [1, 2, 3]. ST technologies come in broadly two flavours, where transcripts are positionally captured from tissue sections via spatially tagged adaptors [4, 5, 6, 7] or via visualization of RNA molecules by *in* situ hybridisation or sequencing, which includes approaches such as *In Situ* Sequencing (ISS) [8, 9], FISSEQ [10], seqFISH [11], MERFISH [12], BaristaSeq [13], STARmap [14], or osmFISH [15]. The former technologies typically cover the whole transcriptome, albeit at a super cellular spatial resolution; whereas the latter IST technologies allow for a subcellular resolution and can cover larger tissue sections, but require one to predefine a set of transcriptomic targets, which are then combinatorially barcoded to enable their parallel sequencing over a few rounds of imaging.

Typical IST data consists of tiled gigapixel cyclic multi-channel images with hundreds of thousands of ‘spots’ of a few pixels in diameter, where each spot corresponds to one RNA molecule whose transcriptomic sequence is fluorescently labeled across different imaging channels and rounds. Decoding of targeted transcripts in order to reconstruct their spatial patterns from such cyclic multi-channel images is therefore an essential task after IST data is acquired. While decoding is conceptually simple, it is complicated in practice by various sources of noise in the data. These include various sources of optical background noise due to tissue autofluorescence or artefact objects, differential illumination, intensity, spectral overlap and diffraction of fluorophores. A further challenge lies in spot detection, which is compounded by microscopic tissue movement between cycles thus requiring stitching and registration such that hundreds of thousands of small spots are well aligned across large images. The resulting distortions and uncertainties, if not properly accounted for, can lead to erroneous decoding.

Many algorithms for analysing IST data have been developed in recent years [16]. Most common schemes for decoding of IST data are based on the argmax approach [8], or its variants [10, 14, 17, 18], where each detected spot is assigned to an encoded sequence (barcode) by selecting the coding channel with the highest image intensity in each round. However, such decoding is sensitive to the aforementioned sources of noise and as a result spots may be assigned incorrectly or to infeasible barcodes, leaving many spots mis-or unlabeled. In particular, different normalisation strategies can greatly alter decoding results since imaging channels may have varied signal-to-noise ratios and thus noise level in one coding channel may be higher than signal in another. An additional source of barcode confusion is the cross-talk between imaging channels because of the spectral overlap of fluorophores. Lastly, some spots may be false positives due to background noise and therefore don’t map to a valid barcode combination of signals.

More recent decoding techniques attenuate some of these issues by computing soft-assignments proportional to distances between barcodes and image values, which are then thresholded [19, 20]. In particular, the approach proposed in [19], firstly transforms barcodes in order to compensate for a cross-talk between coding channels within the same acquisition round, which is estimated based on a *k*-means approach. However, as we demonstrate in this paper, such *k*-means decoding is also susceptible to various types of noise as well as the choice of a threshold value at which barcode assignments are disregarded.

To overcome these shortcomings of existing techniques for decoding IST data, in this paper we introduce a robust decoding method PoSTcode that accounts for different sources of noise and allows for probabilistic assignments of highly variable spot intensities to either targeted or ‘noise’ barcodes. PoSTcode is evaluated against alternative decoding techniques on three large-scale experimental image datasets acquired under different experimental conditions via different versions of the ISS protocol including the latest commercially available Cartana technology (part of 10x Genomics). By comparing the new approach with the state-of-the-art benchmarks, in this paper we show that PoSTcode leads to:

i. increased number of decoded spots,
ii. decreased sensitivity to preprocessing and thresholding of barcode assignments,
iii. decreased confusion of barcodes that differ at one letter only.

Finally, we compare our spot-based decoding against emerging pixel-based approaches such as ISTDECO [21], which are more computationally demanding and harder to aggregate into appropriate spot based signals that are commonly required for downstream analysis such as cell typing [22] or clone mapping [23].

## Results

### IST data acquisition and processing workflow

IST protocols such as ISS acquire signals corresponding to single molecules of mRNA reverse transcribed to cDNA, which bind to molecular probes and are subsequently amplified *in situ* and read out by a set of fluorescent channels *C* > 1 visualized via a microscope. The resulting molecular signals are fluorescent spots of different color depending on the fluorescent channel designed to label a predefined transcriptomic sequence, so that *C* different sequences can be read out simultaneously. Moreover, if such measurements are repeated in multiple sequencing rounds *R* > 1, then combinatorial barcodes can be used for simultaneous sequencing of up to *C*^*R*^ different transcripts. The experimental setting of *in situ* sequencing is schematically illustrated in Fig. 1-a.

**Figure 1:**
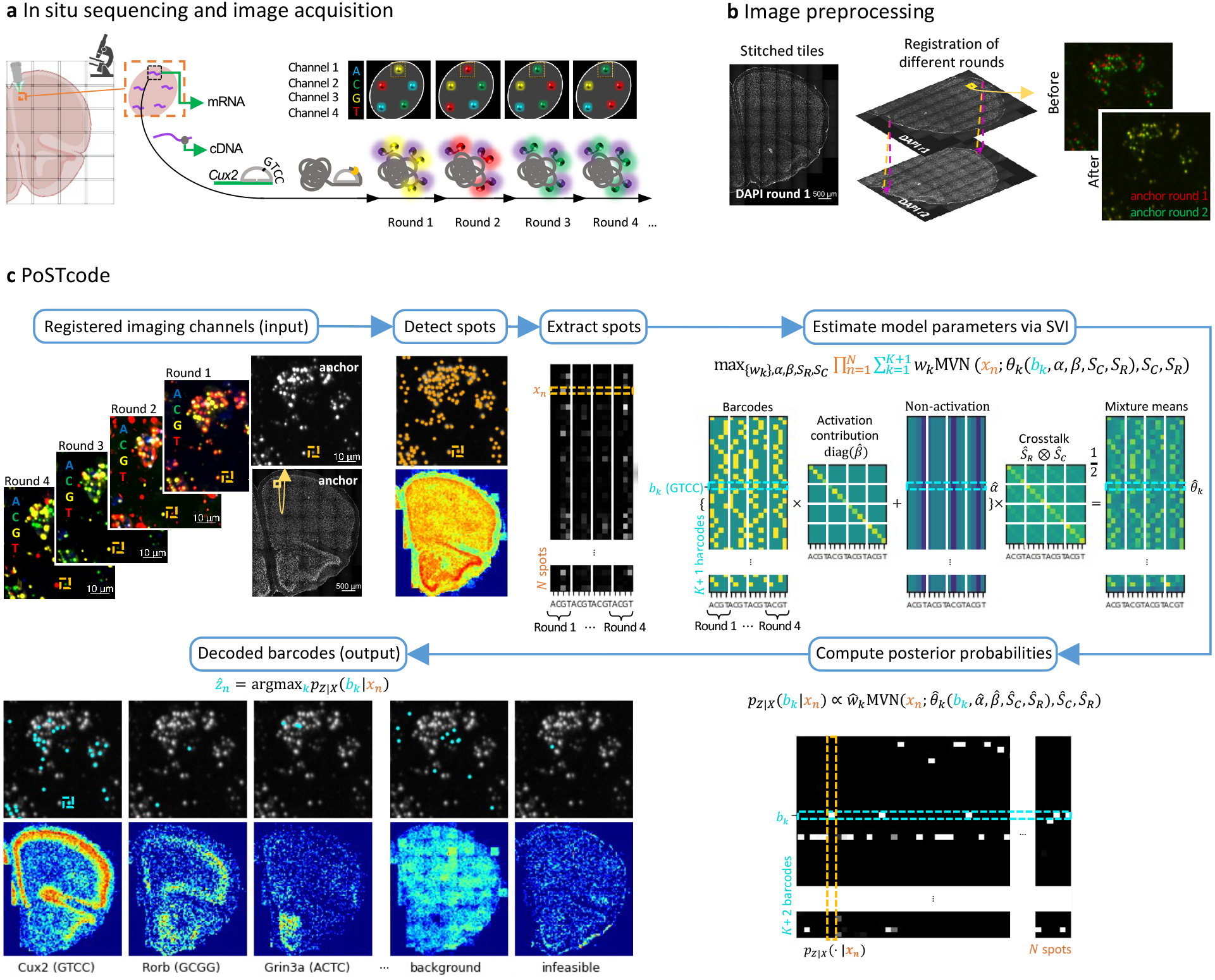
Data generation and processing workflow exemplified on the ISS mouse brain dataset with *K* = 50 barcodes, *R* = 4 rounds, *C* = 4 coding channels and *N* = 394498 detected spots. **a**: Schematic experimental setup. **b**: Image stitching and registration. **c**: Decoding via PoSTcode: Firstly, spot-based signals are detected in the anchor channel leading to spatial locations of all *N* spots. Secondly, values of (top-hat filtered) coding channels are extracted via max-pooling over 3 × 3 pixels area centered at detected spatial locations leading to a data matrix (*x*_1_, …, *x*_*N*_)^T^, whose columns are standardized. Next, model parameters {*w*_*k*_}, *α, β, S*_*R*_, *S*_*C*_ are estimated via SVI per mvGMM given in equations (1)-(2) and posterior probabilities *p*_*kn*_ :=*p*_*Z*|*X*_(*b*_*k*_|*x*_*n*_) per equation (3) are computed. Finally, barcode argmax_*k*_ *p*_*kn*_ is assigned to location *n* ∈ {1,…, *N*} if *p*_*kn*_ ≥ *τ*, for a given threshold parameter *τ* (in all our examples *τ* = 0.7).

Due to a restricted microscopic field of view, approximately 2 × 2 mm^2^ in size, in order to recover a larger field of view across whole tissue, typically of 2 × 2 cm^2^ in size, acquired multi-channel images are firstly stitched in each sequencing round. Additionally, due to movement of a tissue between different sequencing rounds, image registration is required to correct for spatial misalignment of small Gaussian-like spots (approximately 1–1.5 μm in diameter). Corresponding image preprocessing is conceptually illustrated in Fig. 1-b, while algorithmic details of stitching and registration steps are provided in Methods section.

Following image acquisition and preprocessing, in order to reconstruct sequenced transcripts, we propose a decoding procedure PoSTcode, which is illustrated in Fig. 1-c. Similarly to other spot-based decoding techniques for IST data, PoSTcode begins by detecting spots appearing across registered coding channels and extracting intensities from top-hat filtered channels at detected spot locations. Spot detection can be performed via different algorithms such as Trackpy [24] (a Python implementation of Crocker–Grier algorithm [25]) applied to an anchor channel amplifying all targeted transcripts at once, which if not experimentally available can be generated by a suitable projection of coding channels (for more details on spot detection and extraction, see Methods). Once image intensities are extracted from all rounds and channels at *N* detected spot locations, the task is to assign extracted image values {*x*_1_, …, *x*_*N*_} to a collection of barcodes {*b*_1_, …, *b*_*K*_} from a predefined codebook of *K* ≤ *C*^*R*^ elements, which we do via a novel probabilistic model introduced in the following section. Importantly, to make our decoding model robust to variable spot detection sensitivity, and in particular, to false positive detections, we inflate provided codebook by additional ‘background barcode’ *b*_*K*+1_, so that false positive spots extracted from the background noise are assigned to the ‘background’ class, as shown in Fig. 1-c.

### PoSTcode decoding model and estimation of barcode assignments

The driving mechanism behind PoSTcode is a probabilistic model for decoding of extracted spots, which assumes that the stacks of image values extracted from transcript locations are generated by a re-parameterized variant of a matrix-variate Gaussian Mixture Model (mvGMM). Specifically, let {*b*_1_, …, *b*_*K*_} ⊆ {0, 1}^*R*×*C*^ be a codebook of *K* barcodes used in the experiment, which are numerically represented by one-hot encoding of *C* coding channels and *R* sequencing rounds. We model latent labels of observed image intensities by a categorical random variable *Z* taking values in the set of plausible barcodes {*b*_1_, …, *b*_*K*+1_} ⊆ {0, 1}^*R*×*C*^, which includes the ‘noise’ barcode *b*_*K*+1_ consisting of all zeros, according to a prior distribution *p*_*Z*_ parametrised by unknown mixture weights {*w*_1_, …, *w*_*K*+1_} so that *p*_*Z*_(*b*_*k*_) = *w*_*k*_, namely

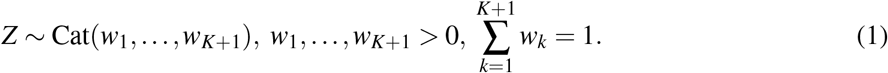

For each barcode *b*_*k*_, *k* = 1, …, *K* + 1, we represent image intensities generated by the corresponding barcoded transcript via a matrix-variate normal random variable defined as

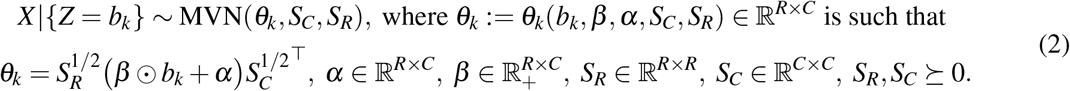

It is important to observe that the mixture mean *θ*_*k*_ is re-parameterized in terms of the scaled barcode *β ⊙ b*_*k*_ + *α*, where the scaling parameters *β* and *α* model channel specific activation and non-activation values, since different channels may have different signal-to-noise ratios. We thereby restrict the search space and instruct the model where to look for clusters in data relative to channel activation points, making it more robust to variable cluster sizes and their shape than a plain mvGMM without such re-parametrization. For example, if there are two channels *c*_*i*_, *i* = 1, 2, in round *r* with their respective non-activation value 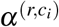 and activation contribution 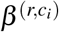, and two barcodes each designed to activate one of the two channels, then our model looks for two clusters of de-correlated image intensities, one around 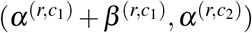 and another around 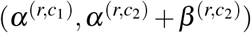, while a plain mixture model looks for two generic clusters. We also note that the model allows for correlation among intensities from different coding channels and different sequencing rounds through covariance matrices *S*_*C*_ and *S*_*R*_.

In practice, we observe a set of *N* image values {*x*_1_, …, *x*_*N*_} ⊆ ℝ^*R*×*C*^ extracted from a stack of *R* × *C* channels at image locations where spots are detected, and we want to decode the set of *N* underlying barcodes that produced those image values. To that end, we assume that *N* desired spot labels are represented by the latent values {*z*_1_, …, *z*_*N*_} ⊆ {*b*_1_, …, *b*_*K*+1_}, which are responsible for the observed values {*x*_1_, …, *x*_*N*_} modelled as the independent observations from the probability distribution 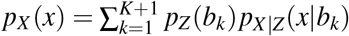 according to the mvGMM specified by equations (1) and (2). We thus decode spot label *z*_*n*_ as the most likely state given observation *x*_*n*_, i.e. as 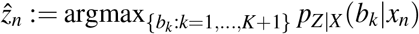, where posterior probabilities

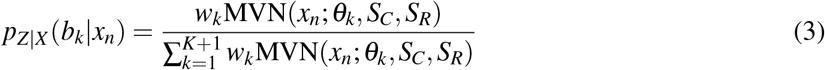

are approximated by plugging in estimated values of mvGMM parameters 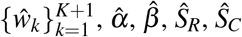. We compute parameter estimates by maximizing the Evidence Lower Bound (ELBO) via the method of Stochastic Variational Inference (SVI) [26, 27] implemented in the probabilistic programming language Pyro [28] (more details available in Methods). In Fig. 1-c, we show the parameters of our model estimated from spots extracted in a mouse brain dataset. In particular, we show the relation of a given codebook to the mvGMM parameters being estimated per equation (2) as well as the resulting posterior values per equation (3).

To increase robustness of our model, after estimating mvGMM parameters, with weight *ŵ*_*K*+2_ = min_*k*=1,…,*K*_*ŵ*_*K*_, we also evaluate the posterior at *b*_*K*+2_ representing any of the ‘infeasible barcodes’ that complement given codebook of *K* ‘feasible barcodes’ to the set of all *C*^*R*^ possible barcode combinations. We find that introducing such a class is useful for removing experimental noise or noise introduced by preprocessing steps such as registration.

Finally, once posterior probabilities per equation (3) are computed, for increased confidence of decoded barcodes, we use a non-trivial threshold value *τ* > 0 to disregard the posterior probabilities (3) that are smaller than *τ*. In our experiments, we found that *τ* = 0.7 works well across different datasets, however we also demonstrate that our decoding results are robust to the choice of hyper-parameter *τ*.

### Model evaluation on three experimental ISS datasets

To illustrate the broad applicability of our decoding algorithm, we evaluate it on three different datasets that were acquired via ISS protocols, each using a different version of the chemistry and being generated on different microscopes. In addition, they each use different numbers of cycles and barcodes and image different tissue types: a mouse brain, a human brain, and a lymph node from a human breast cancer. The total size of their field of view, number of coding cycles *R*, coding channels *C* and barcodes *K*, as well as the total number of spots detected *N*, are reported for each dataset in the table included in Fig. 2. More details about experimental imaging of each dataset and their preprocessing are available in Methods.

**Figure 2:**
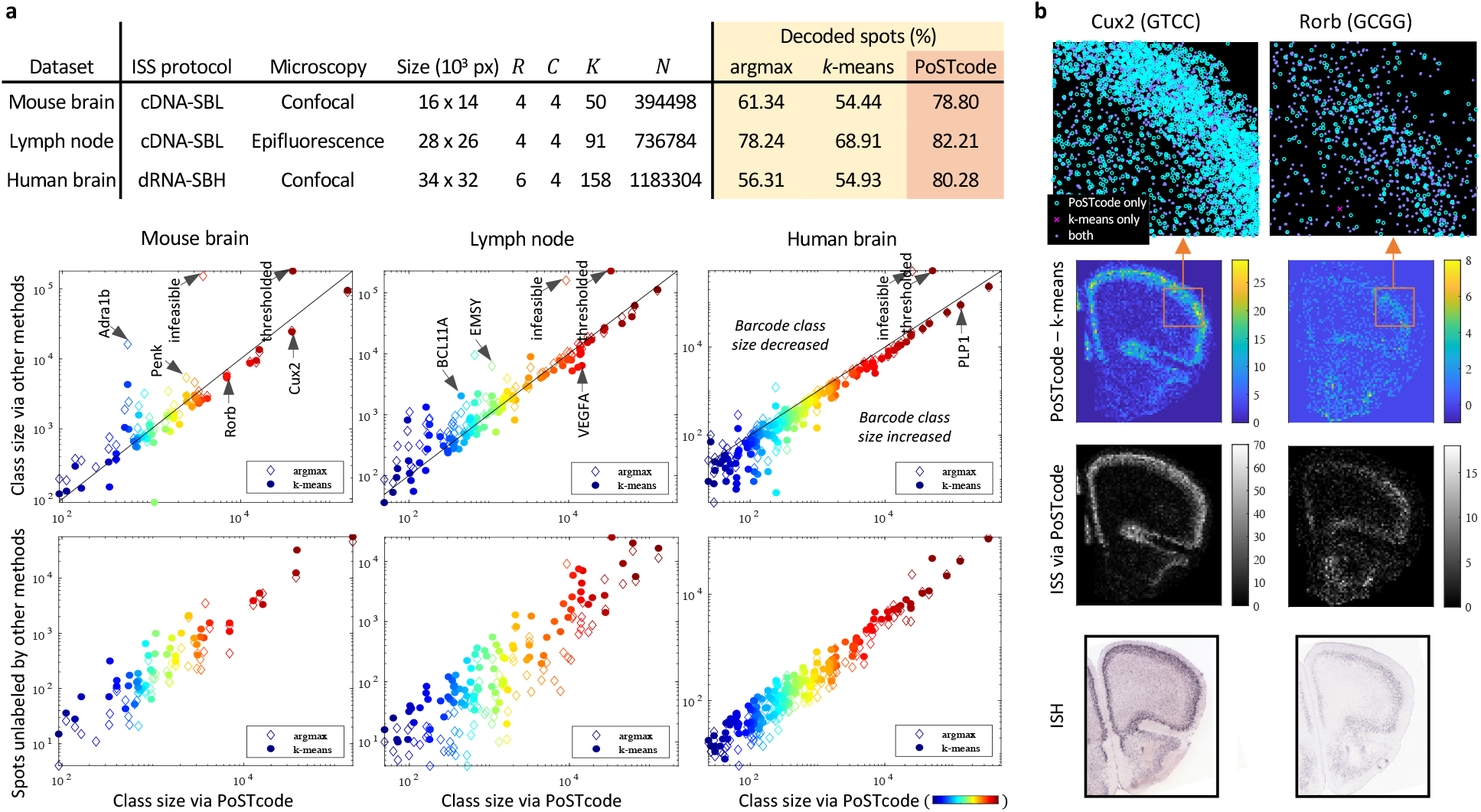
Comparison with alternative methods reveals increased number of decoded spots. **a**: In top row, table of datasets used for comparison with the percentage of decoded spots out of *N* detected spots, which are assigned to *K* encoded barcodes using different decoding methods (dRNA-SBH protocol refers to Cartana technology available from 10x Genomics; for more details on different ISS protocols see Methods). In middle row, size of each barcode class decoded by PoSTcode vs alternative methods in log-log scale. In bottom row, the distribution of spots unlabeled by alternative methods when decoded by PoSTcode (in the argmax/*k*-means case, the ‘infeasible’/’thresholded’ column of the confusion matrix between PoSTcode and argmax/*k*-means, which is included in the SI). **b**: Decoded spatial patterns of two barcodes from the mouse brain dataset, in particular the difference of the decoded signals via two competing methods (with a zoomed section on top), the spatial patterns decoded via PoSTcode, and the anatomically matched sections of the two genes from Allen Mouse Brain Atlas [29].

Even though there is no ground truth, it is possible to visually validate reconstructed spatial patterns of several transcripts by inspecting their spatial expressions obtained in similar tissues via other imaging techniques such as *In Situ* Hybridization (ISH) or Immunohistochemistry (IHC). As it can be seen from Supplementary Fig. 1 and 2, the reconstructed expressions of such transcripts align well with their expected spatial patterns.

In what follows, we demonstrate advantages of our novel probabilistic model by comparing it against alternative decoding models, *k*-means [19] and argmax [8], which are applied to the same set of spots {*x*_1_, …, *x*_*N*_} extracted from locations detected via Trackpy in each dataset. For the comparison purpose, we have implemented the simple argmax approach ourselves, while for the *k*-means we used the original Matlab scripts provided in [19].

### Increased number of decoded spots

In each dataset, PoSTcode’s barcode assignments agree well to those produced by the alternative approaches, however PoSTcode yields more decoded barcodes overall, see Fig 2. By comparing the size of individual barcode classes between the competing methods (middle row of Fig. 2-a), we see that PoSTcode increases mainly well expressed barcodes, while there is a number of smaller barcode classes, particularly in the examples of mouse brain and lymph node, whose size is reduced by PoSTcode. In fact, as we examine in the sections below, the reduction in those classes is because PoSTcode reduces background noise and confusion between barcodes that are close in the Hamming distance.

Since there is no ground truth to compute the accuracy of barcode assignments, we can compare different methods by inspecting most disagreeable barcode classes from the confusion matrices between the competing methods (provided in Supplementary Fig. 1, 2 and 3). Besides disagreement with respect to barcodes that are close in the Hamming distance or sensitive to background noise, the confusion matrices reveal that PoSTcode disagrees with *k*-means/argmax mostly with respect to the thresholded/infeasible (unlabeled) spots, which are mainly assigned to well expressed barcodes when decoded via PoSTcode (bottom row of Fig. 2-a).

For well expressed barcodes with distinct spatial patterns, it is possible to validate that the increased number of decoded spots corresponds to a genuine signal by checking that the difference between the competing methods preserves the expected spatial pattern, as exemplified by the transcripts of *Cux2* and *Rorb* in Fig. 2-b (more examples provided in Supplementary Fig. 1, 2 and 3). In Fig. 2-b, using two well expressed genes characteristic for distinct mouse brain layers of cells, we show that PoSTcode increases them non-uniformly in their respective expected signal regions, therefore enhancing the spatial patterns otherwise produced by *k*-means decoding.

### Increased robustness to preprocessing steps and thresholding

When compared to alternative decoding methods, PoSTcode is more robust to preprocessing noise introduced by imperfect registration (alignment) of spots across different cycles. Correcting misalignment of spots by image registration is an essential but challenging preprocessing step, which can introduce an additional variability of extracted image intensities prior to decoding. In Fig. 3-a and 3-a’, we demonstrate that PoSTcode is significantly more resilient to imperfect registration on the human brain dataset, which is more difficult to correctly register due to the increased number of cycles and removal of the anchor channel from the coding cycles. To correct for a misalignment among the coding cycles, in this dataset we applied a drift correction after initial registration preprocessing and before extracting spots from images (the drift correction is explained Methods). In Fig. 3-a, we show decoded spatial patterns with and without such registration correction of one well expressed barcode with a layer-specific spatial expression, due to which it is easily observable that PoSTcode does not struggle with misregistration as much as other methods. Spatial patterns of additional barcodes with and without drift correction are shown in Supplementary Fig. 4. Moreover, in Fig. 3-a, as a measure of agreement between barcode assignments obtained with or without the registration correction, we show F1 score computed for all barcode classes confirming that PoSTcode barcode assignments with or without such correction disagree overall much less when compared to *k*-means and argmax.

**Figure 3:**
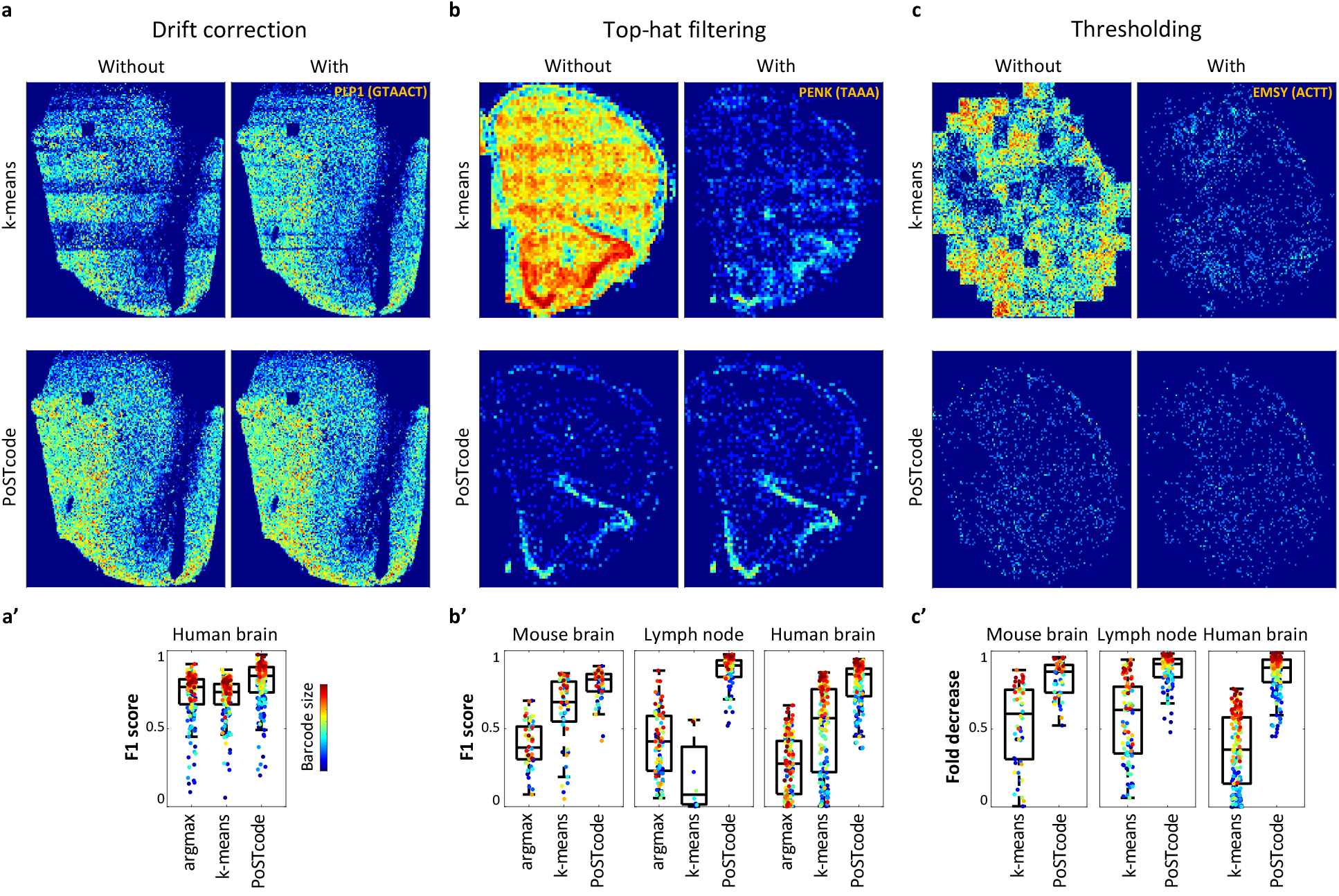
Change in barcode assignments due to imperfect image registration, increased background noise or decreased threshold. **a**: Barcode GTAACT (*PLP1*) from the human brain dataset with and without registration correction decoded by *k*-means or PoSTcode. **a’**: F1 score for each barcode class (different colors ordered from blue to red by increasing class size as in Fig. 2) when decoded using different methods from the spots extracted at the same locations in human brain images before and after registration correction. **b**: Barcode TAAA (*Penk*) from the mouse brain dataset with and without top-hat filtering of background noise decoded by *k*-means or PoSTcode. **b’**: F1 score for each barcode class with and witouth top-hat filtering for three different datasets. **c**: Barcode ACTT (*EMSY*) from the lymph node dataset with recommended threshold (0.9 for *k*-means and 0.7 for PoSTcode) and the zero threshold. **c’**: Fold decrease in each barcode class when threshold is changed from 0 to the recommended value.

Beside spot-like signals, experimental images often contain additional background noise due to autoflorescence or other optical artifacts, which increases variability of values extracted from imaging channels. Such background noise is typically decreased prior to decoding by applying top-hat filtering with prescribed diameter of spot-based signal that should remain in images. In Fig. 3-b and 3-b’, we demonstrate that PoST-code is significantly more robust to background noise than alternative methods, by comparing the decoding results with and without top-hat filtering. In particular, in Fig. 3-b, we show spatial patterns of a barcode class that is highly sensitive to background noise (images of additional barcodes are shown in Supplementary Fig. 4), while in Fig. 3-b’, we compare the F1 scores of decoded spots with and with top-hat filtering for all barcode classes decoded via different methods. From the overall higher F1 scores, we can see that PoSTcode’s assignments are much more consistent regardless of whether optical noise is present or not. By inspecting the spatial patterns of barcode classes that are highly sensitive to top-hat filtering, such as *Penk* shown in Fig. 3-b, we also observe that, unlike PoSTcode, alternative decoding methods substantially reduce such classes if top-hat filtering is applied. However, even when top-hat filtering is applied, such classes are still inflated and unreliably decoded via other methods, while PoSTcode produces expected spatial patterns with or without top-hat filtering.

Both PoSTcode and *k*-means decoding require a tuning parameter to be chosen for the final decoding step where small scores of barcode assignments are thresholded and dismissed as non-reliable. In Fig. 3-c and 3-c’, we demonstrate that PoSTcode is more robust to the choice of its threshold value by comparing the reduction (fold decrease) in each barcode class when threshold value is increased from zero to the respective recommend value (0.7 for PoSTcode and 0.9 for *k*-means). More generally, we also confirm that PoSTcode’s threshold does not affect its decoding results as much as it does for *k*-means, by computing the distance between the distribution of barcode assignments when different values of threshold are used, see Supplementary Fig. 4-c and 4-d. This is partially achieved by including ‘noise’ barcode classes, but more importantly, by increased confidence in barcode assignments demonstrated by overall lower entropy (Supplementary Fig. 4-b).

### Decreased barcode confusion

In the lymph node and mouse brain datasets, each with four coding rounds, there is a significant number of barcodes that differ in one round only, i.e. barcodes that are at the Hamming distance one (211 and 59 pairs respectively), which are more likely to be mistaken for each other thereby producing correlated spatial patterns. By closely inspecting the confusion matrices between different decoding methods (Supplementary Fig. 1 and 2), the disagreement of PoSTcode with respect to the alternative decoding methods is the highest for such barcode pairs. Specifically, in the lymph node example, the highest values in the confusion matrices are for the pairs (*BCL11a* (CGGT), *VEGFA* (CGAT)) and (*VAV1* (TTTA), *CD24* (TTAA)), while in the example of the mouse brain, for barcode pairs (*Adra1b* (AACG), *Gapdh* (TACG)) and (*Sstr1* (GACC), *Cux2* (GTCC)). In Fig. 4-a, we show one example of these pairs (others are shown Supplementary Fig. 5), where it can be seen that their corresponding spatial patterns are much more correlated when decoded by alternative approaches than when decoded by PoSTcode.

**Figure 4:**
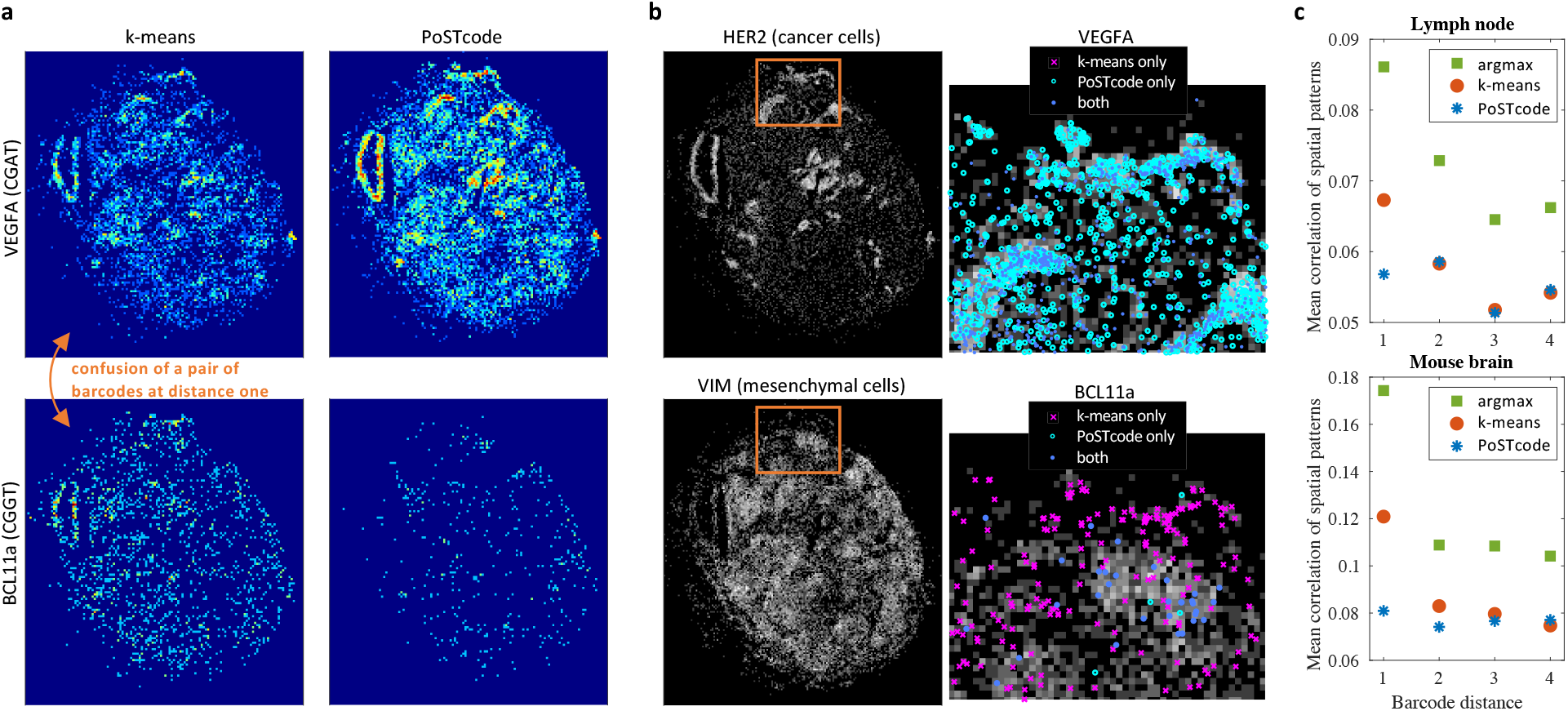
Decreased confusion of barcodes at the Hamming distance one. **a**: Spatial patterns of barcodes CGAT (*VEGFA*) and CGGT (*BCL11a*) decoded by competing methods. **b**: Spots decoded as *VEGFA* / *BCL11a* against a grey heatmap of spots decoded as gene *HER2* / *VIM* (characteristic for cancer / mesenchymal cells). **c**: Mean correlation among the spatial patterns of barcodes against their Hamming distance for two different datasets.

Specifically in the example shown in Fig. 4-a, we know that gene *BCL11a* is normally expressed in germinal center B-cells, while *VEGFA* is normally increased in hypoxic regions of cancer aggregates particularly in areas marked by necrosis. However, we can see that when decoded by *k*-means, *VEGFA*, associated with cancer aggregates, bleeds into *BCL11a*, which is not expected in high abundance among dense cancer regions. In Fig. 4-b, locations of *BCL11a* are shown against a grey heat-map expression of VIM associated with mesenchymal cells, where it is more likely to be found, while locations of *VEGFA* are shown against a grey heat-map expression of *HER2* associated with cancer cells, where *BCL11a* is unlikely to be found. This reveals that the spatial pattern of *BCL11a* obtained by *k*-means is indeed more diffused across the dense cancerous regions, where *VEGFA* is expected.

In Fig. 4, we also investigate how correlation between spatial patterns is related to the Hamming distance of the underlying barcodes by plotting the average correlation of spatial patterns between all pairs of barcodes against their Hamming distance. Even though there is no reason for any particular trend between the two factors, when using the alternative decoding methods, in both datasets the correlation between spatial patterns increases when the barcode distance is equal to one. PoSTcode on the other hand does not show an increase in the correlation between close barcodes.

By reducing barcode confusion, PoSTcode allows more flexibility in barcode design. Although in principle, barcode confusion can be minimized by using barcodes that differ in at least two rounds, such constraint increases the number of rounds needed to accommodate a codebook of a given size. Enabling accurate distinction of barcodes that differ in a single sequencing round allows larger number of barcodes with a given number of rounds, or the same number of barcodes with a smaller number of rounds.

### Comparison with pixel-based decoding

Recently two methods for pixel-based decoding of IST data have been proposed, ISTDECO [21] and BarDensor [30]. These methods start from registered (background-removed) images and in order to circumvent running spot detection algorithms on such images, they directly recover a large non-negative matrix with regression coefficients assigning pixels to barcodes, which can then be thresholded and aggregated into coordinates of decoded spots. As such, these methods are more computationally demanding, and also require precise pixel-level registration as well as fine-tuned spatial deconvolution and aggregation into spot-like signals. The central promise of pixel-based approaches is to detect more spots in crowded areas with many close signals. Yet we note that close spots may also be detected by Trackpy or related spot detection algorithms if the parameter controlling the minimal separation between spot centers is decreased.

The two pixel-based approaches are compared in [21], out of which we used the faster and the more recent one, ISTDECO, to compare against our spot-based decoding procedure PoSTcode, see Fig. 5. For a fair comparison, we ran ISTDECO with parameters fine-tuned by the authors on the ISS dataset used in their paper and originally analyzed in [19], which consists of 180 tiles of size 512 × 512 pixels. We found it to be 18.75 times slower than our decoding procedure, which takes 5.09 min including running time of Trackpy’s spot detection (78% of total PoSTcode running time). Even though ISTDECO uses the crosstalk compensated barcodes as its input, which the authors estimated by running *k*-means decoding on spots detected in Matlab as in [19], we did not measure that additional computational time.

**Figure 5:**
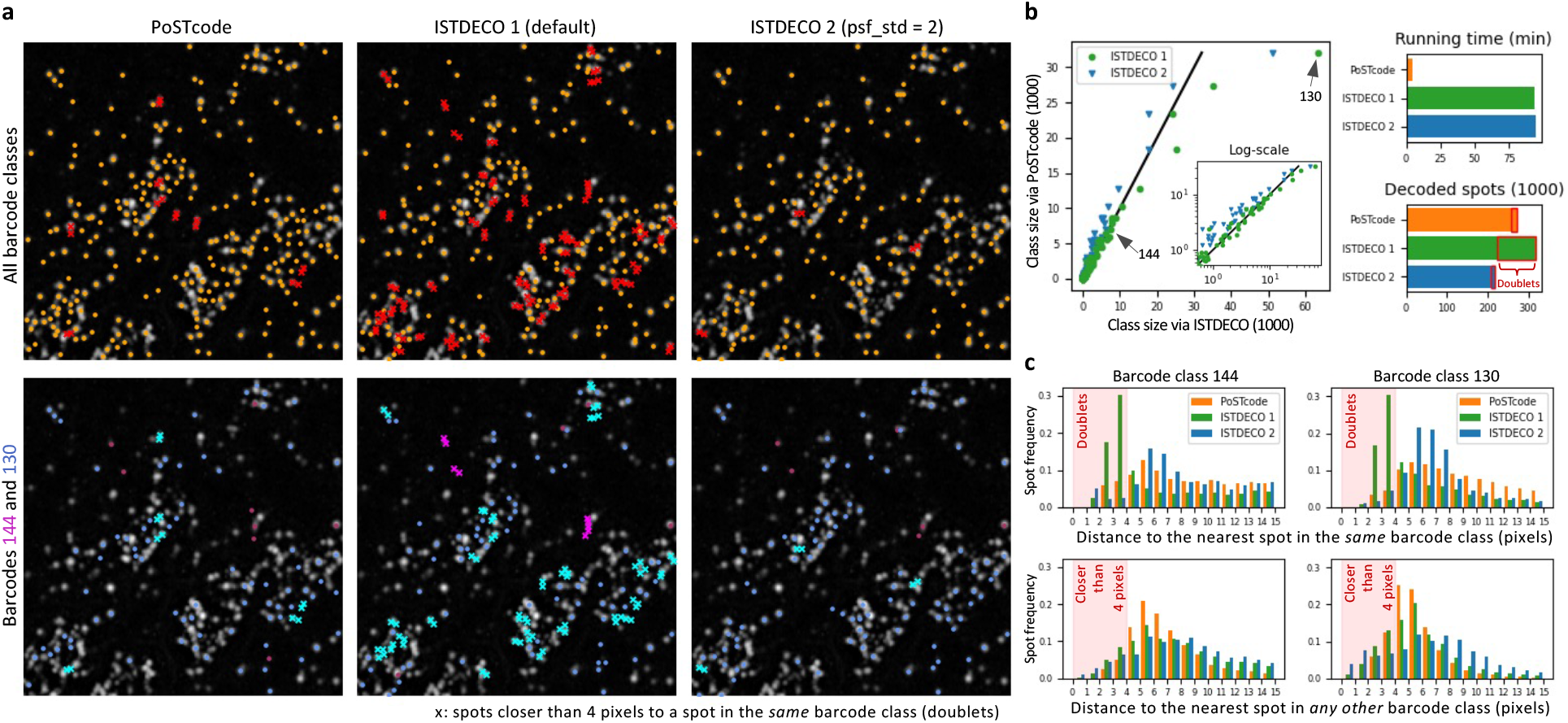
Comparison of PoSTcode and pixel-based decoding ISTDECO, where ISTDECO 1 denotes the use of the default value of parameter psf std = 1 while ISTDECO 2 denotes the use of psf std = 2. **a**: Top row shows all decoded spots in a crowded FOV, where doublets (defined as spots within the same barcode class that are closer than 4 pixels) are marked in red crosses; while the bottom row shows two different barcode classes (doublets of barcode 144 in magenta and doublets of barcode 130 in cyan). **b**: Sizes of barcode classes when decoded via different methods as well as the total number of decoded spots and algorithmic running time. **c**: Histograms of all spots from barcode classes 144 and 130 grouped based on their distances to the nearest spot in the same barcode class (top) or to the nearest spot from any other class (bottom).

When compared to PoSTcode’s decoded coordinates that were detected via Trackpy (with spot diameter 5 px, minimum separation between spots 2 px, percentile of spots removed 0), ISTDECO with its default parameters seems to decode 14% more spots overall, out of which more than 70% are assigned to the most prevalent class (denoted as barcode 130 in Fig. 5-b). However, as it can be seen from images shown in Fig. 5-a, ISTDECO’s coordinates within the same barcode class are often clustered together indicating that a single signal source is often split into multiple counts, likely due to an inappropriate spatial deconvolution. In fact, unlike PoSTcode, ISTDECO’s empirical distribution of distances to the closest spot within the same class is sharply peaked at less than 5 pixels, which is a typical spot diameter, see Fig. 5-c. Yet this effect is restricted to the same barcode class, and unlike the wider empirical distribution of distances to the closest spot from any other class, indicating that it results from erroneously splitting a signal rather than true molecular crowding. In fact, when removing these likely doublets, PoSTcode decodes a greater number of singleton transcripts.

The problem of multiplet counts could potentially be alleviated by increasing ISTDECO’s tuning parameter ‘psf std’, which corresponds to the standard deviation of the Gaussian shaped point-spread function used for spatial deconvolution (by default set to 1), to indicate a larger spot size. Yet in this case the total number of spots decoded by ISTDECO drops below the number decoded by PoSTcode (psf std = 1.5, 8% less; psf std = 2, 20% less). Furthermore, the most prevalent class stays overly inflated when decoded via ISTDECO.

## Discussion

In this paper we introduced PoSTcode, a Bayesian matrix variate Gaussian mixture model for decoding imaging spatial transcriptomics data in the presence of different sources of experimental and optical noise. When compared to existing methods for decoding barcodes sequenced via image-based spatial transcriptomics, we showed that PoSTcode is more reliable across different datasets while at the same time it enhances spatial patterns of decoded transcripts and reduces their mislabeling. Although for the purposes of this paper we used three image datasets selected to illustrate different aspects of the proposed approach against the existing decoding methods, PoSTcode has been further applied to dozens of image datasets acquired at two different labs, namely at Wellcome Sanger Institute and ScilifeLab [23].

Although PoSTcode has been specifically tuned for the experimental setting of *in situ* sequencing where barcodes are designed so that exactly one imaging channel is active per sequencing round, the method introduced here is not restricted to such barcode design. While not in the focus of this study, PoSTcode may be extended for decoding combinatorial codebooks as utilised in protocols such as MERFISH [12, 20] typically decoded by a pixel-based approach.

Even though we were able to validate PoSTcode’s favourable performance on multiple large scale datasets, there are still avenues for further improvement. Specifically, PoSTcode requires the preprocessing steps of image registration and spot detection that can introduce noise. In particular, although PoSTcode achieves higher stability to mis-registration, the underlying decoding model operates on the assumption that spot locations are properly aligned across different sequencing rounds. As we have seen in the human brain example, after initial image registration, alignment of spots can be improved by using Trackpy to track detected spots and correct the average drift between different imaging rounds. However, such spot tracking requires additional parameter fine-tuning, so it remains to be seen if more accurate alignment can be learnt from the data in a more automated way.

Furthermore, spot detection algorithms can be overly sensitive to optical noise thereby selecting spots that are not produced by actual transcripts, which is especially pronounced if spot detection sensitivity is increased. PoSTcode accounts for such noise by encoding it with additional barcodes, namely by introducing the background class during estimation of model parameters and by evaluating the posterior approximation at infeasible barcodes. Even though by considering ‘noise’ barcode classes, the noise introduced by overly sensitive spot detection is reduced, it may be computationally demanding to add all possible (*C* + 1)^*R*^ ‘noise’ classes during estimation and properly estimate their corresponding weights.

Beside the parameter that controls their sensitivity, classical spot detection algorithms such as Trackpy require additional input parameters such as spot size and minimal spot separation, which may be variable across different images and thus require fine-tuning. Pixel-based approaches attempt to address some of these problems by avoiding applying spot detectors on noisy images. However, they still rely on a fine-tuned spatial deconvolution and post-decoding aggregation, which as we have seen, may produce unsatisfactory results.

Future work may therefore focus on automated decoding starting from raw data and involve simultaneous registration, detection and decoding of spots based on their combinatorial encoding with minimal fine-tuning. One avenue towards this would be to train a deep neural network on images reliably decoded with fine-tuned registration and spot detection, so that it can automatically account for noise introduced by these preprocessing steps.

## Methods

### Experimental framework

In Results section, PoSTcode is evaluated using three different experimental datasets: a mouse brain, a human lymph node and a human brain. We sequenced the mouse and human brain samples at Welcome Sanger Institute, while the lymph node sample was sequenced in ScilifeLab from Stockholm, Sweden, and was previously used for the analysis reported in [23]. In what follows, we provide details of the experimental framework used for sequencing these samples.

### Tissue specimens

Fresh frozen healthy human primary motor cortex (age 57yr, male) was obtained from Edinburgh Brain Bank (HMDMC: 19/0120). Cryosections (10 μm) were collected on a cryostat (Leica), mounted onto superfrost glass slides (VWR) and stored at -80°C. For fresh frozen mouse brain section, a wild-type adult C57BL/6 mouse (postnatal day 68, female) was transcardially perfused with ice-cold phosphate-buffered saline (PBS) and heparin. The brain was then embedded in OCT (Tissue-Tek) and frozen on isopentane-dry ice slurry and stored at -80°C. Coronal cryosections (10 μm) were collected on a cryostat (Leica), mounted onto superfrost glass slides and stored at -80°C. Mouse brain sample processing was carried out at the Wellcome Sanger Institute in accordance with UK Home Office regulations and UK Animals (Scientific Procedures) Act of 1986 under a UK Home Office license, which were reviewed by the institutional Animal Welfare and Ethical Review Body. For human lymph node, serial 10um tissue sections were cut from fresh frozen tumour tissue blocks mounted on superfrost slides and sent from DFCI to ScilifeLab where ISS and IHC was performed under Karolinska Institutes rules for the handling of blood and other human sample material, reference number 1-31/2019 (with a local HUMRA risk assessment form).

### Target selection and Padlock probe (PLPs) design

For mouse brain, mouse neocortex single cell RNA-seq data from [31] and mfishtools R library [32] were utilized to select 50-plex marker genes to delineate neuron and glia cell types in mouse brain. For human human brain, 158-plex CNS probe design was carried out by CARTANA (Sweden) using similar methods as in [19] and also commercially available as follows: Neuro General [N1C v1.0], CNS Excitatory [N2D v1.0], CNS Glia [N3E v1.0] and CNS Inhibitory [N4F v1.0]. For human lymph node, 91-plex oncology gene panel design was carried out as a part of [23]. The panel include genes related to proliferation, EMT, invasiveness, stemness, angiogenesis as well as genes for breast cancer subtyping and oncotypeDX recurrence scoring.

#### *In situ* Sequencing (ISS)

Two different ISS protocols, cDNA-SBL (Sequencing By Ligation) and dRNA-SBH (Sequencing by Hybridization), have been utilized for the manuscript. For cDNA-SBL (mouse brain and lymph node), after tissue fixation (4% PFA for 30 min at RT) and permeabilization (0.1M HCl with 0.1 mg/ml pepsin (Sigma) for 90 s 37°C), SecureSeal reaction chambers were mounted on top of the tissues (Grace Biolabs) and cDNA was synthesized *in situ* using specific DNA primers (5nM) and random decamer primers (5μM) (IDT). Rnase H was used to generate single-stranded cDNA that the padlock probes could hybridize to. Hybridized padlock probes (10 nM) (IDT) were ligated using Tth ligase, a highly specific DNA ligase that can discriminate correct base-pairing at the single nucleotide level. Only completely target-complementary padlock probes become ligated, forming closed circles that could then be amplified through rolling circle amplification (RCA). The four nucleotide target-specific barcodes included in the padlock probe linker sequence were clonally amplified in the RCA products allowing identification through sequencing by ligation using anchor probe and fluorophore-labelled interrogation probes [8]. Nuclei were stained with 4’,6-diamidino-2-phenylindole (DAPI). The target-specific barcodes were sequenced with four sequencing and imaging rounds.

For dRNA-SBH (human brain), 10x Genomics provided reagents in kits (High-Sensitivity library preparation kit) with an accompanying protocol that was followed. Briefly, after tissue fixation (3.7% Formaldhyde for 30 min at RT) and permeabilization (0.1M HCl for 5 min at RT), probe mix was incubated on tissue section overnight in hybridization buffer followed by stringent washes and then incubated in a ligation mix. After washes, RCA was performed overnight and labelled for detection. Nuclei were stained with DAPI. The target-specific barcodes were sequenced with six sequencing and imaging rounds.

### Image acquisition

For human and mouse brain, cyclic images were acquired with a Perkin Elmer Opera Phenix High-Content Screening System, in confocal mode using a *z*-stack of 0.8-1μm separated 20-21 slices and a tile overlap of 7%. Images were scanned with a 20x objective. For each tile, four different coding channels were imaged corresponding to four different bases, for human brain: A–AF488, G-AF647, C-Atto425 and T-AF568, and for mouse brain: A-Atto 425, G-Atto 565, C-AF488 and T-AF647. Additionally, all image tiles included a DAPI channel used for nuclei staining, while for the mouse brain example, they also included anchor channel Atto490LS to locate all sequenced transcripts at once. For the first base sequenced, the exposure times were calibrated so that the signal intensity values were similar for all sequencing channels, and the calibrated exposure times were then kept constant for all remaining sequencing cycles. For human lymph node, images were acquired with an automated Zeiss Axioplan II epifluorescence microscope (Zeiss, Oberkochen, Germany) using a *z*-stack of 0.49μm separated 11 slices and a tile overlap of 10%. All image tiles included six channels (Nucleus-DAPI, A-Cy5, G-Cy3, C-Texas Red, T-AF488, and anchor-Cy7) and were scanned with a 20x objective.

### Computational framework

The computational framework used for image pre-processing and decoding is presented below in greater detail. All algorithmic step starting from acquired image data are listed in Pseudo-code 1. We emphasize that when compared to other methods, PoSTcode improves decoding results due to a novel probabilistic model used to assign barcodes to image values extracted at detected locations of RNA signal. However, prior to spot decoding, RNA signals need to be detected and corresponding image values extracted from registered and top-hat filtered coding channels. Additionally, due to a restricted microscopic field of view, one may want to stitch multi-channel images in each round in order to recover a larger filed of view. These pre-processing steps can be performed using different algorithms, and in what follows, we describe how each of these steps was performed in all our experiments.

### Image stitching and registration

In all microscopic tiles, we integrated *z*-axis via the max-projection, which was followed by stitching along remaining spatial axes *x* and *y*. Stitching of microscopic tiles was performed separately in each round using DAPI channel. For the mouse and human brain dataset, exported tiles were stitched using Acapella scripts provided by Perkin Elmer., while for the lymph node dataset, tiles were stitched using built-in stitching processing method of ZEN Blue version 2.3SP1 (Carl Zeiss Microscopy, Jena, Germany).

All datasets were then registered in two steps. Firstly, we used feature registration algorithm implemented in Python via OpenCV-contrib library (version 4.3.0) [33] to compute an affine transformation of DAPI channel from round *r* > 1 (moving image) with respect to DAPI channel from the first round *r* = 1 (reference image). In particular, key points were detected using the FAST feature detector, whose surrounding areas were described using the DAISY feature descriptor, while the FLANN-based

**Pseudo-code 1:**
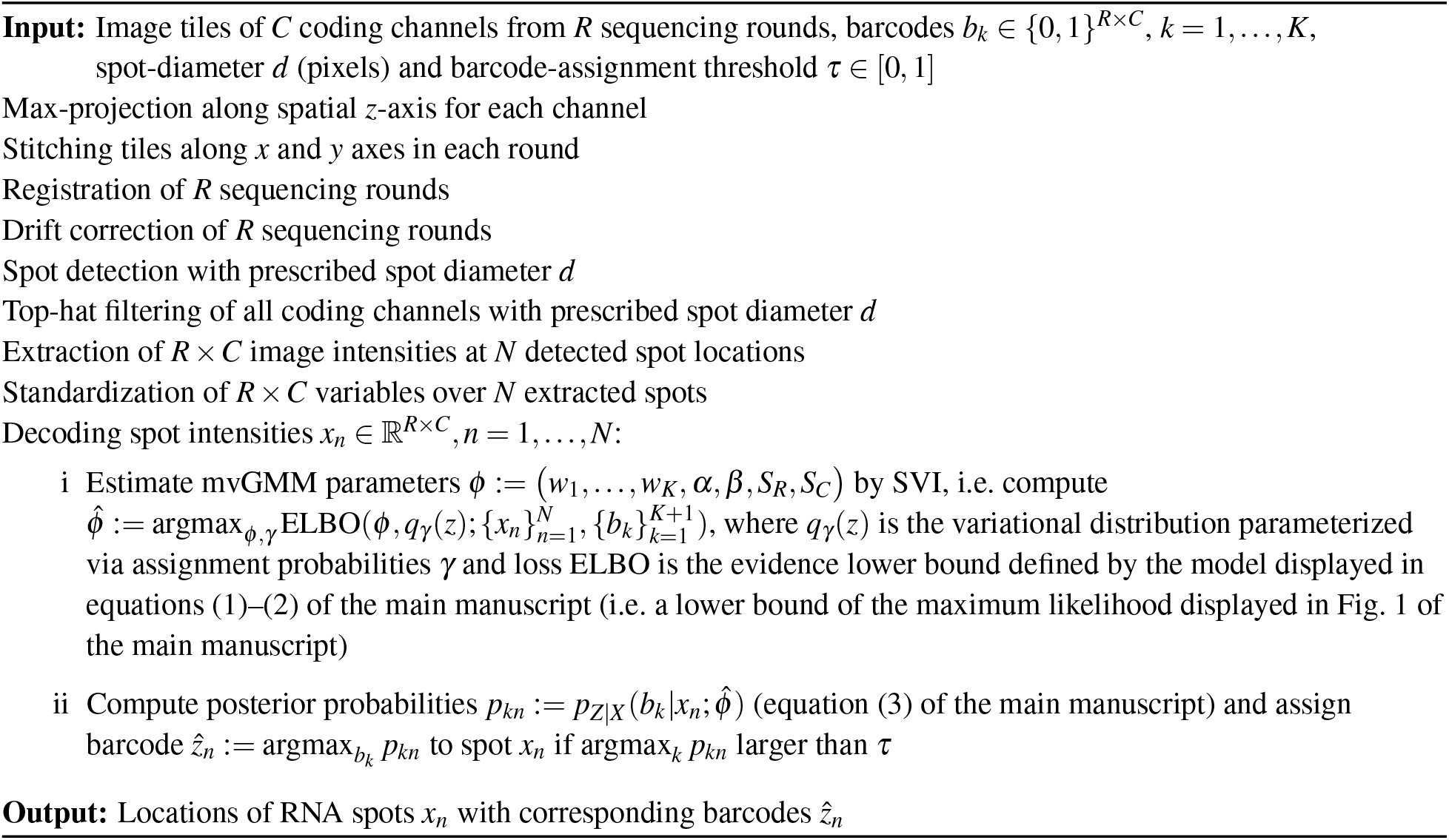
Main algorithmic steps starting from experimentally acquired images

matcher was used to find correspondences between pairs of key points from reference and moving images and filter out unreliable points. Lastly, the remaining key points were processed using the RANSAC-based algorithm that aligns them and estimates affine transformation parameters with four degrees of freedom. For the second registration step, a nonlinear registration algorithm based on Farneback optical-flow available in Python via OpenCV library was used to achieve more accurate registration by warping images locally. Specifically, local warping was computed using anchor channel (mouse brain and lymph node), or DAPI channel (human brain), from round *r* > 1 with respect to the corresponding channel of the first round. The computational pipeline implementing these registration steps was optimized so that it can be performed efficiently on large images and the corresponding code for feature registration is available at github.com/BayraktarLab/feature_reg, while the code for optical-flow registration at github.com/BayraktarLab/opt_flow_reg.

### Drift correction

Additionally, in the human brain example, registration quality was further improved by applying a linear drift correction to tiled images after the two registration steps outlined above. Drift correction was implemented via publicly available software Trackpy [24], which can be used to track the spots across different rounds. Specifically, the spots were first detected in each round via Trackpy’s function ‘trackpy.batch’ applied to the anchor channel in each round. Since the human brain dataset doesn’t have experimentally obtained anchor channel, for this step a fake anchor was created by the max-projection of all normalized coding channels from the corresponding round, where coding channels were normalized by their respective 95th percentile. Spots were subsequently linked via ‘trackpy.link df’ in order to compute the median displacement of spots from round *r* > 1 with respect to the first round *r* = 1 along both spatial axes. In case the median displacement was larger then zero, the sequencing round was shifted in the corresponding displacement direction in order to improve spatial alignment of spots with respect to the first round.

### Spot detection and extraction

Gaussian-like spots were detected via Trackpy [24] with a given spot diameter *d*. For all examples used in this paper, *d* was set to 5 pixels. This step can be performed either by linking detected spots in each round via Trackpy’s functions ‘trackpy.batch’ and ‘trackpy.link df’, or provided satisfactory registration results and not overly noisy anchors, simply by locating spot coordinates using only one of the rounds via ‘trackpy.locate’. The later approach is preferred since it allows detection of spots in overly crowded areas by decreasing the default minimal separation between any two spot centers (from *d* + 1 to 2 pixels). In particular, in the mouse and human brain examples, we used the later approach, while in the lymph node example, we detected spots by the former approach via spot linking since the anchor channel there was overly noisy and linking helped removing noise. Spot detection was applied on tiled images of the experimentally acquired anchor channel (mouse brain and lymph node) or the fake anchor (human brain), which was obtained by the max-projection of normalized coding channels from the first sequencing round.

Following spot detection, we extracted corresponding image intensities from top-hat filtered coding channels via max-pooling over 3 × 3 pixel area centred at detected spot coordinates. Top-hat filtering was applied with a disk of diameter *d* = 5 in order to decrease background noise, while max-pooling was applied to increase robustness to miss-alignment of spots. All the algorithmic steps listed up to here were applied prior to decoding via any of the spot-based decoding methods used for benchmarking.

### Spot decoding via PoSTcode

Prior to estimation of model parameters, *N* extracted spots {*x*_1_, …, *x*_*N*_} ∈ ℝ^*R*×*C*^ were standardized to achieve a Gaussian like distribution of image values in each coding channel as it is a common practice when applying GMMs. Specifically, extracted data was standardized so that values from each channel have mean zero and standard deviation one after log-transformation is applied to the channel values. In particular, for each channel *c* ∈ {1,…,*C*} and round *r* ∈ {1,…, *R*}, transformation 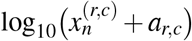 was firstly applied to the extracted image values 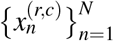, where constant 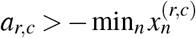 is computed so that the 60th-percentileof 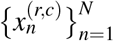 is mapped to the middle of the log-transformed interval. Following this log-transformation, each channel (*r, c*) was standardized by subtracting the channel mean and dividing by its corresponding standard deviation.

Parameters of the model defined by equations (1)–(2) were estimated using SVI [26, 27] available via the probabilistic programming language Pyro [28], which is built on top of the deep learning framework PyTorch thereby allowing for automatic differentiation as well as utilization of GPUs if available. Specifically, we used ‘pyro.infer.SVI’ with a model that was implemented following equations (1)–(2) in their vectorized form, and a guide function (variational distribution) that was automaticly generated via ‘pyro.infer.autoguide.AutoDelta’ applied to the model function. In addition, ELBO loss function ‘pyro.infer.TraceEnum ELBO’ was used in order to enumerate over discrete latent variables, which was maximized for model parameters using Adam stochastic gradient descent [34] available through ‘pyro.optim.Adam’. To make the model scalable for large *N*, the log likelihood term of the ELBO was computed using mini-batch sub-sampling.

Although the most likely states can be directly computed via ‘pyro.infer.infer discrete’ after training the model, we instead implemented E-step of the EM algorithm [35] to compute the assignment probabilities via equation (3) by plugging in parameter estimates. This step then allows computation of assignments to additional barcodes that were not included in the estimation procedure described above. The additional barcodes used in this E-step correspond to ‘infeasible’ barcodes, which were not targeted in the experiment but may appear in the data due to misalignment/optical noise. For such ‘infeasible’ barcodes, as well as for the ‘background’ barcode, we used a small value of the corresponding mixture weight *ŵ*_*k*_, *k > K*, which is set to the minimal value of the weights estimated using ‘feasible’ (targeted) barcodes min_*k*=1,…,*K*_ *ŵ*_*k*_.

## Code and data availability

PoSTcode is available at github.com/gerstung-lab/postcode, which includes a notebook demonstrating how to decode an example tile from the mouse brain dataset available in the same git repository, an example tile available through the starfish package, and a notebook for reproducing the mouse brain decoding results presented in the paper. The associated mouse brain dataset is available at www.ebi.ac.uk/biostudies/studies/S-BSST700.

## Author contributions

M.Ga. and M.Ge. designed the decoding methodology; M.Ga. developed and validated the method, analysed decoding results, and drafted the manuscript; J.S.P., O.B. and M.Ge. provided feedback on the decoding results and the manuscript; J.S.P., T.L., V.V. and O.B. contributed to the design of the computational and experimental workflow required prior to decoding; J.S.P., K.R., J.S., C.S. generated ISS data; M.N. and L.R.Y. designed ISS experiments; J.S.P., T.L. and V.V. registered images. All authors reviewed and commented on the manuscript.

## Acknowledgments

This work was supported by OpenTargets grant OTAR2070. We acknowledge the *in situ* sequencing unit, part of the spatial and single cell biology platform, funded by SciLifeLab for technical assistance with the ISS experiments. We would also like to thank Artem Lomakin and Jose Guilherme Coelho Peres de Almeida for comments on a preprint of this manuscript.

