## Supplementary Figures for "PoSTcode: Probabilistic image-based spatial transcriptomics decoder"

### Supplementary information for
### "PoSTcode: Probabilistic image-based spatial transcriptomics decoder"

Milana Gataric[1], Jun Sung Park[1,2], Tong Li[2], Vasyl Vaskivskyi[2], Jessica Svedlund[3], Carina Strell[3,4], Kenny Roberts[2], Mats Nilsson[3], Lucy R. Yates[2], Omer Bayraktar[2], and Moritz Gerstung[1,5]

[1]European Molecular Biology Laboratory, European Bioinformatics Institute EMBL-EBI, Hinxton, UK
[2]Wellcome Sanger Institute, Hinxton, UK
[3]Science for Life Laboratory, Department of Biochemistry and Biophysics, Stockholm University, Solna, Sweden
[4]Department of Immunology Genetics, and Pathology, Uppsala University, Uppsala, Sweden
[5]Division for AI in Oncology, German Cancer Research Centre DKFZ, Heidelberg, Germany

#### 1 Supplementary note

To complement Fig. 2 of the main manuscript, in Supplementary Fig. 1, 2 and 3 for each of the datasets (mouse brain, lymph node and human brain respectively) we show complete confusion matrices (in the log-scale) between PoSTcode and each of the alternative decoding methods (argmax or $k$-means), along with several barcodes with distinct, layer-specific spatial patterns whose spots were largely unlabeled when alternative methods were used (assigned to infeasible class by argmax or thresholded by $k$-means). For such barcodes, by checking the spatial pattern of the difference between the competing methods (PoSTcode − argmax or PoSTcode − $k$-means), we can see that the signal enhanced via PoSTcode corresponds to the expected spatial patterns originally decoded via any of the three methods. Moreover, for further visual evaluation of the signals enhanced via PoSTcode, in the mouse brain example, we also show ISH data of the genes that were available from the Allen Brain Atlas ISH database repository, and in the lymph node example, we show IHC for *HER2* gene.

To complement Fig. 3 of the main manuscript, in Supplementary Fig. 4-a, we show additional examples of barcodes from the human brain dataset demonstrating robustness of PoSTcode to different pre/post-processing steps. In addition, in Supplementary Fig. 4-b, we compute entropy of barcode assignments averaged over all spots assigned to the same barcode class. Since $k$-means scores take values between 0 and

1, but not necessarily sum up to one, in order to compute their entropy, we first normalized the scores for each spot with their $\ell_1$-norm. Much lower entropy of PoSTcode's assignments indicates increased confidence in assigning barcodes to spots and thus likely stability to noise and the choice of threshold value. Furthermore, in Supplementary Fig. 4-c and d, we confirm stability of PoSTcode to thresholding by showing that the histograms of barcode assignments at different threshold values do not differ a lot and as much they do for *k*-means. In fact, from these plots we can see that the choice of threshold value for *k*-means is very important since we can obtain very different results with similar threshold values.

Finally, to complement Fig. 4 of the main manuscript, in Supplementary Fig. 5, we show additional pairs of barcodes that differ at one letter only, which correspond to the highest entries in the confusion matrices displayed in Fig. 1 and 2 (highest other than the entries of the columns with spots unlabeled via alternative methods).

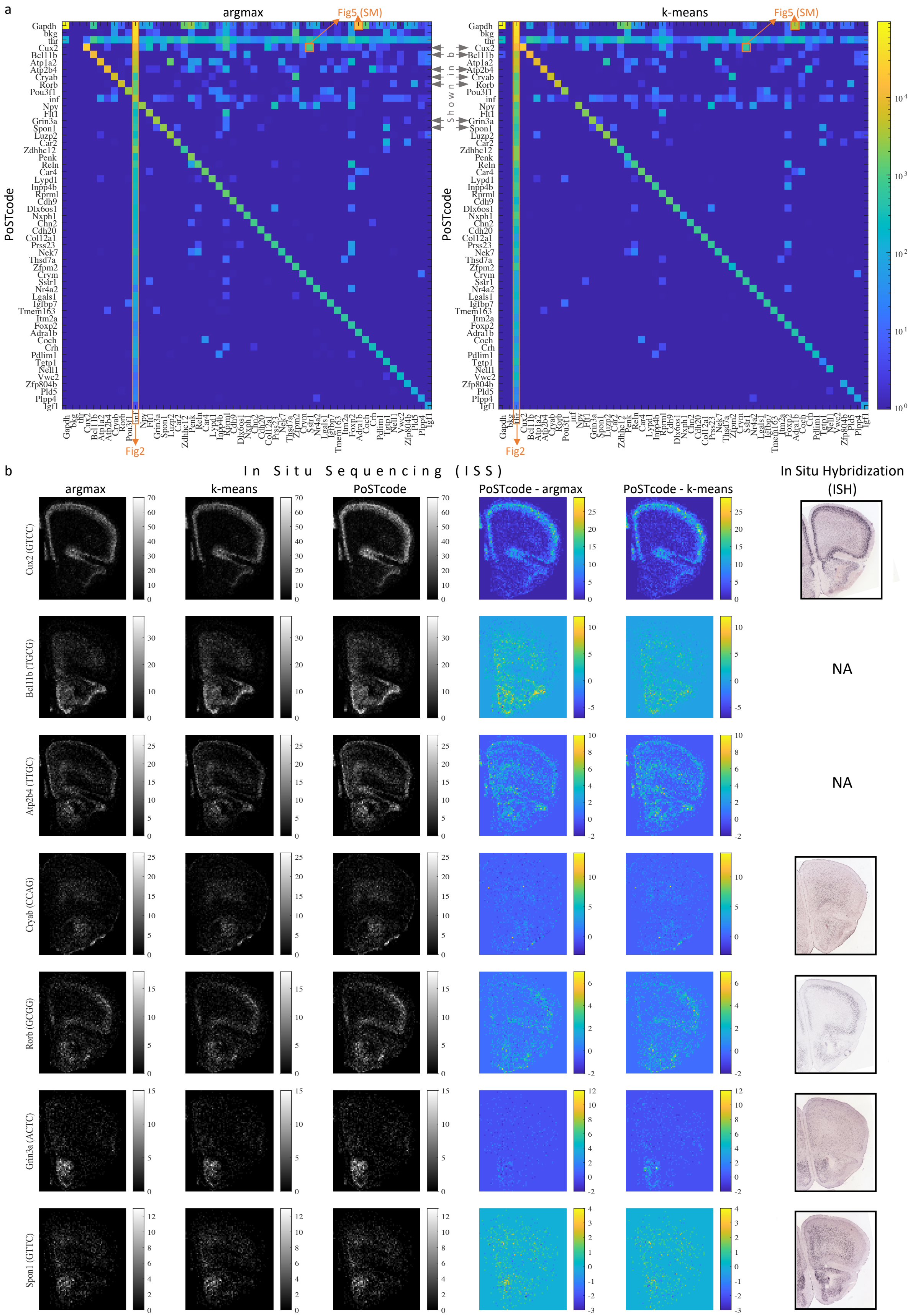

**Supplementary Figure 1:** Complementing Fig. 2 of the main manuscript for the Mouse brain example.

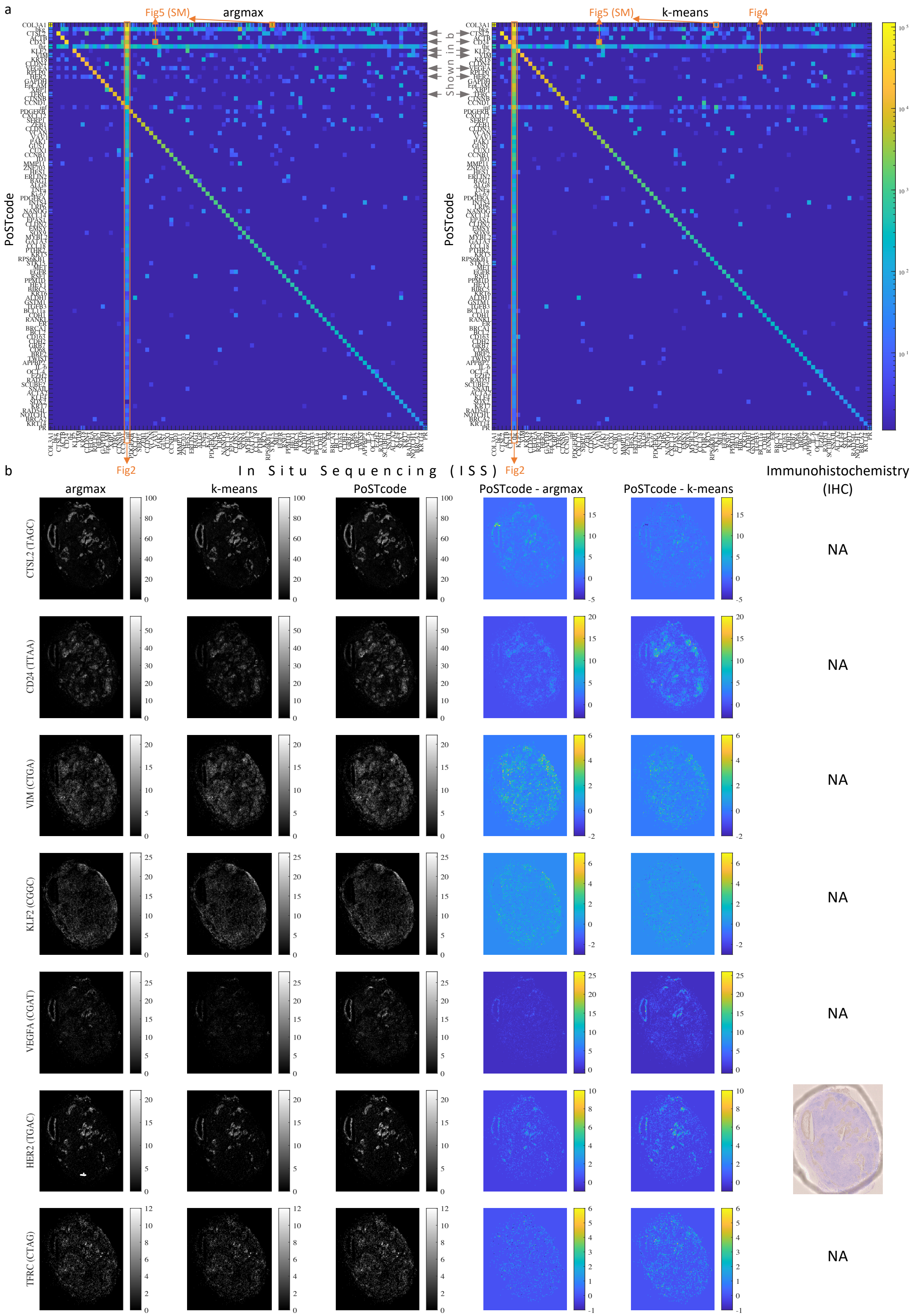

**Supplementary Figure 2:** Complementing Fig. 2 of the main manuscript for the Lymph node example.

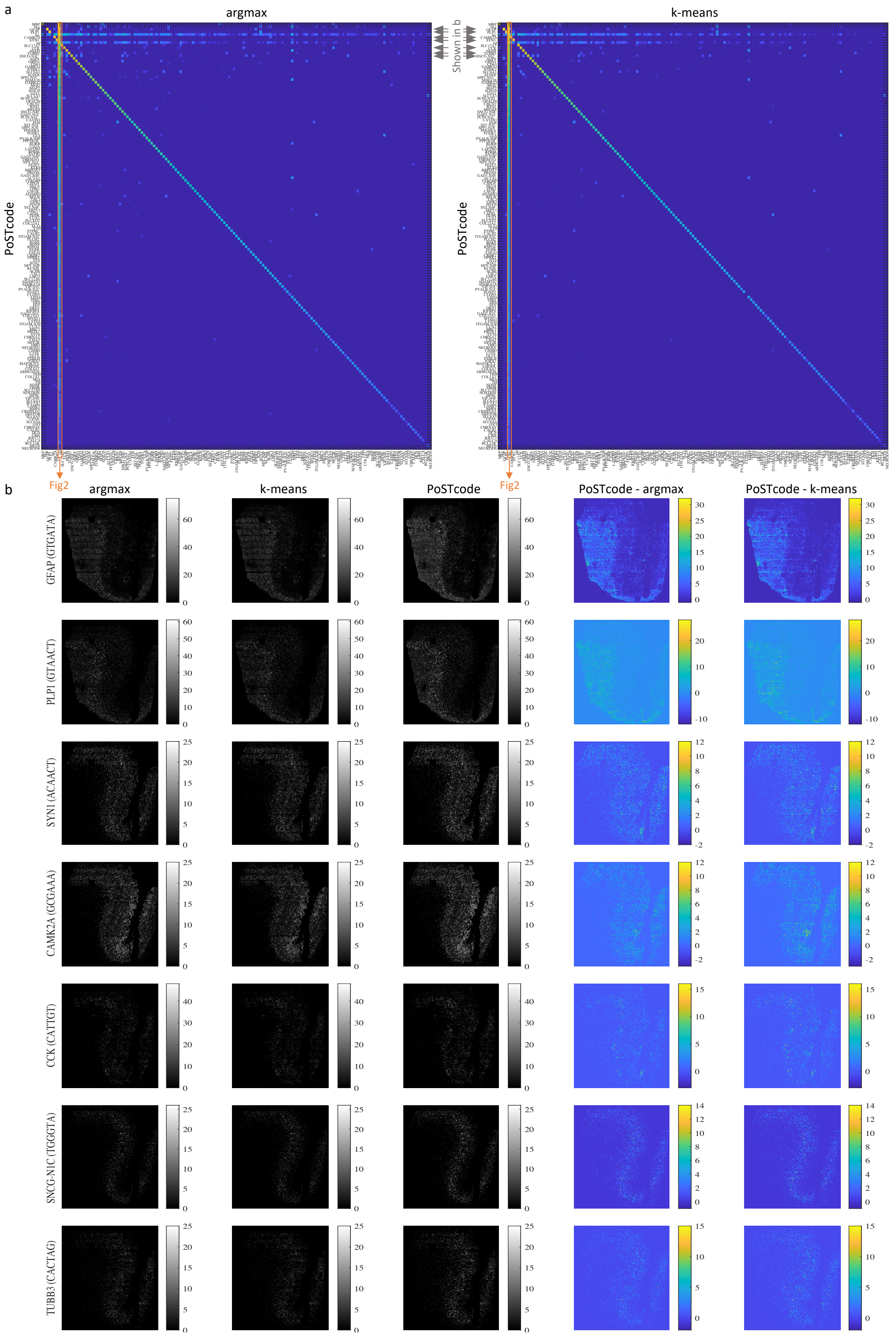

**Supplementary Figure 3:** Complementing Fig. 2 of the main manuscript for the Human brain example.

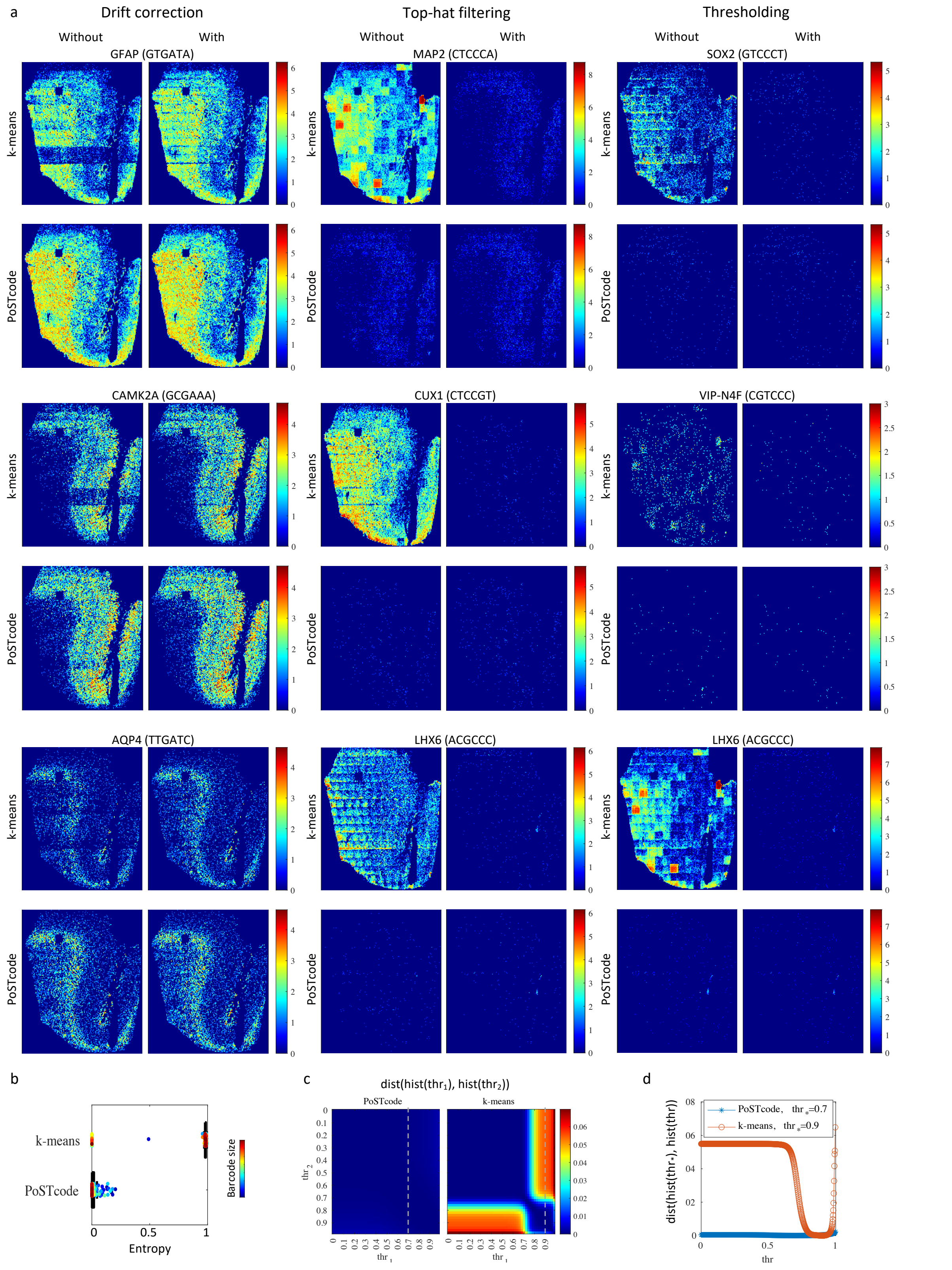

**Supplementary Figure 4:** **a**: Additional examples demonstrating robustness of PoSTcode (images are shown in log-scale and so that each group of four is on the same scale). **b**: Mean entropy of barcode assignments where the mean is take over all the spots with the same most likely barcode class. **c**: Distance between two histograms of barcode frequencies obtained at two threshold values measured as $1 - \cos$ of the angle between the histograms. **d**: 1D intersection of the plots from **c** where one of the threshold values is fixed to the recommended value of the corresponding method.

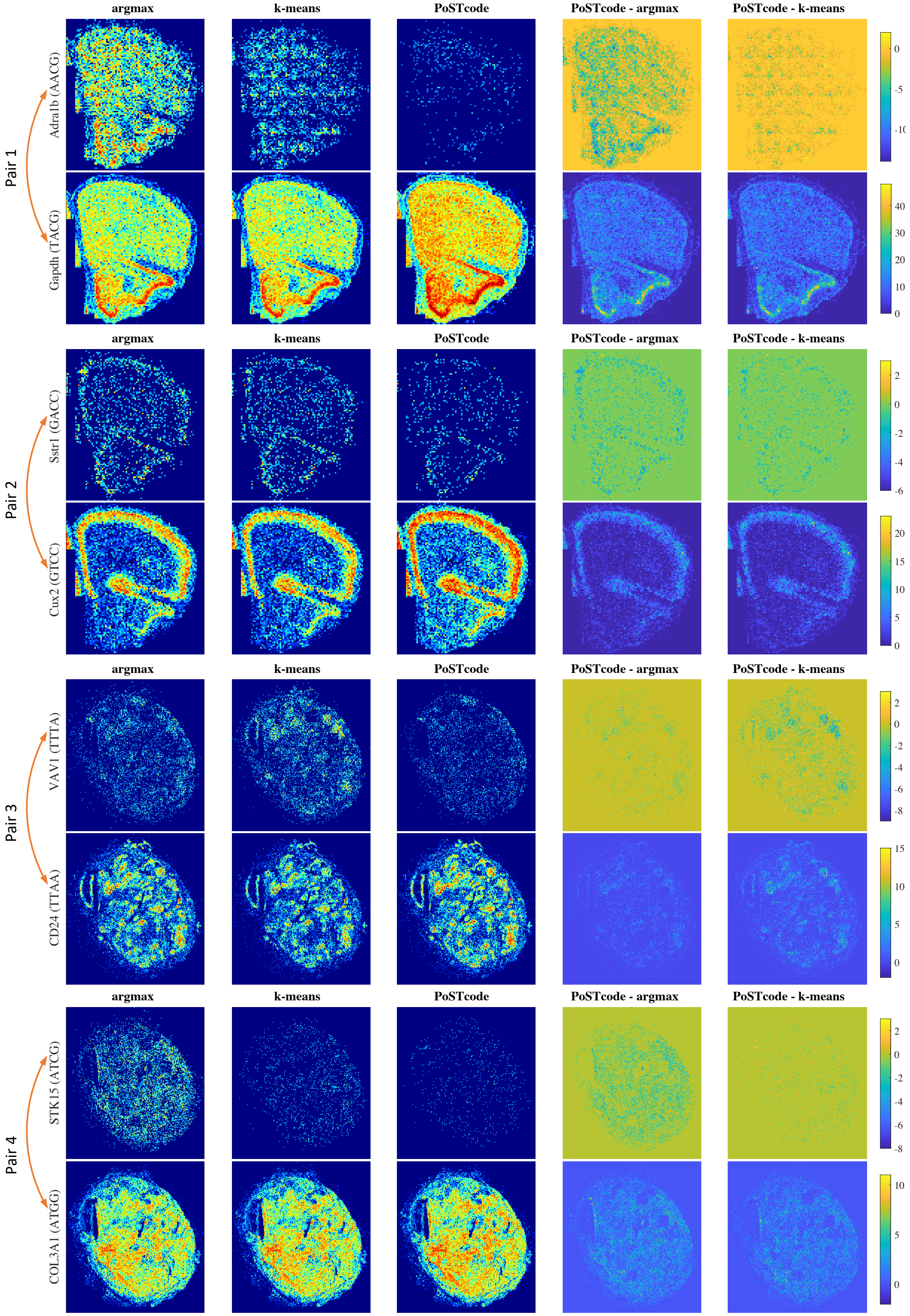

**Supplementary Figure 5:** Additional examples of confused barcode pairs.
